# Naturally occurring N-terminal mutations in SARS-CoV-2 nsp1 impact innate immune modulation but do not affect virus virulence

**DOI:** 10.1101/2024.11.23.624996

**Authors:** Ruchi Rani, Mohammed Nooruzzaman, Leonardo C. Caserta, Diego G. Diel

## Abstract

The non-structural protein 1 (nsp1) of SARS-CoV-2 plays a key role in host innate immune evasion. We identified two deletion variants (Δ82-85 and Δ83-86) in the N-terminal region of the nsp1 of a SARS-CoV-2 BA.5.2.1 variant recovered from a human patient. Analysis of the sequence databases revealed a frequency of 0.5% of these mutations amongst available SARS-CoV-2 sequences. Structural analysis of the deletion mutant nsp1Δ82-85 and nsp1Δ83-86 revealed a distortion in the protein pocket when compared to the wild-type nsp1 which may affect protein function. To evaluate the functional relevance of these mutations, we cloned the mutant BA.5.2.1 nsp1Δ82-85 and nsp1Δ83-86 and wild-type nsp1 proteins in expression plasmids and performed luciferase reporter-based assays to assess activation of the interferon and nuclear factor kappa B (NF-κB) signalling pathways. Both nsp1Δ82-85 and nsp1Δ83-86 mutants showed marked decreased ability to inhibit the interferon beta (IFN-β) and NF-κB pathway activation. To assess the relevance of these deletions in the context of SARS-CoV-2 infection, we generated recombinant viruses carrying the wild type BA.5.2.1 nsp1 or the BA.5.2.1 nsp1Δ82-85 and nsp1Δ83-86 deletions in the backbone of WA1 strain. *In vitro* characterization of the recombinant SARS-CoV-2 viruses revealed that the recombinant viruses containing the nsp1Δ82-85 and nsp1Δ83-86 deletions presented similar plaque size and morphology to those produced by the wild-type rWA1-BA.5.2.1-nsp1 virus, indicating a similar ability of the mutant viruses to spread from cell to cell. Importantly, pathogenesis studies revealed that these mutations did not affect virus virulence and pathogenesis in a hamster model of SARS-CoV-2 infection.

## Introduction

Severe acute respiratory syndrome coronavirus 2 (SARS-CoV-2) is the causative agent of coronavirus disease 19 (COVID-19) which resulted in an unprecedented pandemic (Wu et al., 2020). SARS-CoV-2 is an enveloped single-stranded positive-sense RNA (+ssRNA) virus that belongs to the genus *Betacoronavirus* within the family *Coronaviridae* (Xu et al., 2020). Due to its RNA nature, the virus rapidly evolved accumulating genetic diversity while circulating in the human population (van Dorp et al., 2020). The accumulation of mutations has the potential to impact the ability of the virus to modulate host immune responses as well as its ability to cause disease and spread in susceptible population. This highlights the importance of monitoring SARS-CoV-2 genetic diversity and of characterizing the functional effect of identified mutations on virus infection and pathogenesis (Akaishi et al., 2022; Arya et al., 2023; Chen et al., 2021; Jeronimo et al., 2024; McCarthy et al., 2021; Yuan et al., 2021)

The SARS-CoV-2 nsp1 is functions as the host shutoff factor that is expressed after cell entry to repress multiple steps of host protein expression (Kamitani et al., 2006)(Thoms et al., 2020). The N-terminus of nsp1 binds to the 40S ribosomal subunit causing the degradation of host mRNAs and thus inhibiting the synthesis of host proteins, while the C-terminus of nsp1 plays an important role in stabilizing the ribosomal interaction with the protein (Schubert et al., 2020). Additionally, nsp1 impacts the host immune response by downregulating the expression of interferon-stimulated genes (ISGs) (Kumar et al., 2021). Moreover, nsp1 also interferes with host mRNA export from the nucleus to the cytoplasm, dampening host antiviral innate immune responses. These mechanisms enable the virus to replicate and are thought to contribute to virus virulence and pathogenesis (Zhang et al., 2021).

Here we characterized two nsp1 N-terminal deletion mutations (Δ82-85 and Δ83-86) that we identified during the genome surveillance of the SARS-CoV-2 Omicron BA.5.2.1 variant. We determined the effects of deletion mutations (nsp1-BA.5.2.1Δ82-85 and nsp1-BA.5.2.1Δ83-86) on modulation of host innate immune response *in vitro*, and further assessed their effect on virus virulence and pathogenesis using a hamster model of SARS-CoV-2 infection.

## Materials and Methods

### 3D homology model generation

We identified two four amino acid deletions (nsp1Δ82-85 and nsp1Δ83-86) in the N-terminus of the nsp1 protein of a SARS-CoV-2 BA.5.2.1 variant isolated from a human patient (NYI20/22). The sequences of the variant were retrieved and subjected to structural analysis using the Swiss-model homology tool, using PDB ID 8A55 as the template for modeling. Both the target sequence and template (.pdb) files were uploaded to the SWISS-MODEL server, and 3D structure models were generated for both mutants and wild type BA.5.2.1 sequences. These models were then analyzed for structural features and validated using a Ramachandran plot via the PROCHECK server. The root mean square deviation (RMSD) of the models was also calculated using the PyMOL Molecular Graphics System, Version 2.3.4, Schrodinger, LLC (Delano, 2002; *PyMOL* | *Schrödinger*).

### Luciferase reporter assay

The nsp1 gene of SARS-CoV-2 variants B.1 and BA.5.2.1 wild-type and two variants with deletion mutations (nsp1Δ82-85 and nsp1Δ83-86) were cloned into the mammalian expression plasmid pLVX-EF1a-IRES-Puro, in-frame with an N-terminal Strep II tag. The deletion nsp1Δ82-85 and nsp1Δ83-86 variants were amplified by RT-PCR from a clinical isolate, cloned into the pLVX-EF1a-IRES-Puro plasmid, and the resultant constructs were transformed into *E. coli* Stable cells (NEB® Stable Competent *E. coli*) for amplification.

For the luciferase assays, HEK293T cells were seeded in 24-well cell culture plates at a density of 2 × 10□ cells/well in complete minimum essential media (MEM) (Gibco™, Waltham, MA, USA) supplemented with 10% fetal bovine serum (FBS) and cultured at 37°C in a 5% CO□ atmosphere. After 24 hours, the medium was replaced with fresh complete MEM containing 10% FBS, and the plate was returned to the incubator. Transfection was performed using Lipofectamine 3000 reagents (Invitrogen, USA) with pIFN-β-Luc, pNF-κB-Luc or pIRF3-Luc (200 ng/well) and pRN-Luc (50 ng) reporter plasmids in combination with 250 ng/well of wild-type (WT) nsp1, the deletion nsp1Δ82-85 and nsp1Δ83-86 variants, or an empty pLVX-EF1a-IRES-Puro vector (EV) as a control. After 24-hour incubation, the cells were stimulated with Sendai virus (SV, Cantell strain) (100 hemagglutination unit/well) or TNF-α (25 ng/well), and incubated for 12 hours. After incubation, the media was removed, and the cells were lysed in 50 µL of 1X Passive Lysis Buffer (Promega, USA). The plate was then incubated at room temperature (RT) for 10 minutes with continuous shaking. Following incubation, the plate was transferred to -80°C to ensure complete dissociation of the cell membrane. The cell lysates were then thoroughly mixed by scraping the wells with pipette tips. Next, 10 µL of the cell lysate was transferred to a white 96-well MicroWell Greiner Microlon luminometer plate. Luciferase activity was measured using the dual luciferase assay according to the manufacturer’s instructions (Dual-Glo® Luciferase Assay System, Promega, USA). For the firefly luciferase assay, 50 µL of Dual-Glo luciferase reagent was added to each well, and the plate was incubated for 10 minutes at RT in the dark before measuring firefly luminescence using the luminometer (BioTek Synergy LX Multimode Reader). Subsequently, 50 µL of Dual-Glo Stop & Glo reagent was added to each well, followed by another 10-minute incubation at RT in the dark. The ratio of luminescence obtained from the target reporters (pIFN-β-Luc, NF-κB-Luc or IRF3-Luc) to luminescence from the control Renilla reporter (pRN-Luc) was calculated to normalize the transfection efficiency. Then, the relative IFN-β-, NF-κB- or IRF3-driven luciferase activity was calculated as fold change over the unstimulated cells as described elsewhere (Martins et al., 2024).

### Western blot for luciferase activity assay

To examine the expression of nsp1 protein in luciferase reporter assays, Western blotting was performed. HEK293T cells (2×10^5^ cells/well) seeded in 24-well plate were transfected in triplicate with pIFN-β-Luc, pNF-κB-Luc or pIRF3-Luc (200 ng/well) and pRN-Luc (50 ng) reporter plasmids in combination with 250 ng/well of wild-type (WT) nsp1, the deletion nsp1Δ82-85 and nsp1Δ83-86 variants, or an empty pLVX-EF1a-IRES-Puro vector (EV) as a control. After 24 hours, cells were stimulated with SeV (100 HAU) or TNF-α (25 ng/well) as described above for 12 hours or kept unstimulated. Stimulated and unstimulated samples of pIFN-β-Luc, pNF-κB-Luc, and pIRF-3-Luc transfected cells were separated on a 10% SDS-PAGE gel using the Mini-PROTEAN® Tetra Vertical Electrophoresis System (Bio-Rad, USA). Following electrophoresis, proteins were transferred onto nitrocellulose membranes (TarnsBlot® Turbo™ Midi-Size Nitrocellulose, Bio-Rad, USA) using the Trans-Blot® Turbo™ Transfer system (Bio-Rad, USA) for 7 minutes. Furthermore, the membrane was blocked with 5% non-fat dry milk powder (Research Product International, USA) in tris buffer saline containing 0.2% Tween-20 (0.2% TBST) buffer overnight at 4°C to prevent non-specific binding. After washing three times with 0.2% TBST buffer for 5 min at room temperature, the membranes were incubated with an anti-Strep tag II antibody (THE™ NWSHPQFEK Tag Antibody, GenScript, USA, 1:1000 dilution) to detect Strep-tagged nsp1 and with an anti-GAPDH mouse-antibody (1:1000 dilution, Santa Cruz Biotechnology, Inc., USA) to serve as a loading control. Subsequently, the respective secondary antibody was added at a 1:10,000 dilution. A protein molecular weight marker (10–250 kDa, Dual Color Protein Standard Marker, Bio-Rad) was used for determining the protein sizes. The images were captured and analyzed using a ChemiDoC™ MP Imaging System (Bio-Rad, USA).

### Generation and characterization and functional analysis of rSARS-CoV-2 virus

To assess the relevance of the nsp1Δ82-85 and nsp1Δ83-86 deletions in the context of SARS-CoV-2 infection, recombinant viruses were generated, carrying the wild-type BA.5.2.1 or the nsp1-BA.5.2.1Δ82-85 and nsp1-BA.5.2.1Δ83-86 deletions, within the backbone of the WA1 strain cloned into the pBeloBAC backbone; kindly provided by Dr. Luis Martinez-Sobrido. The cloning process for the recombinant wild-type (nsp1-BA.5.2.1) and variant viruses (nsp1-BA.5.2.1Δ82-85 and nsp1-BA.5.2.1Δ83-86) involved PCR amplification of two key fragments: the pBeloBAC backbone (ranging from the *ApaI* restriction site up to the start of the nsp1 sequence, including 20-nucleotide sequence) and the nsp1 region (beginning after the 20-nucleotide region of the pBeloBAC backbone and extending through the nsp1 sequences up to *KasI* restriction enzyme site). The differences between the wild-type and variants are located near the *KasI* restriction enzyme site that is incorporated at the time of PCR amplification of nsp1 region. Further, Gibson assembly (ThermoFisher Scientific, USA) was used to ligate the fragments, and the ends of the PCR-amplified products included *ApaI* (in the pBeloBAC backbone) and *KasI* (in the nsp1 region) restriction sites. Both enzymes were utilized for digesting the pBeloBAC-WA1 backbone and PCR products, followed by overnight ligation. The ligated constructs were then electroporated into ElectroMax™ DH10B™ cells (Invitrogen, USA). After electroporation, cells were incubated at 37°C for 2 hours, centrifuged at 1000 rpm for 5 minutes, and plated on chloramphenicol plates, which were subsequently incubated at 37°C. After 24-48 hours, colonies were picked and confirmed by sequencing.

The confirmed constructs were then transfected into Vero E6 TMPRSS2 cells using the Lipofectamine 3000 reagents (ThermoFisher Scientific, USA), and the P0 virus was rescued within 96 hours. Subsequent passages were carried out up to the P3 stage, after which sequence identity and integrity of the recombinant viruses was verified by whole genome sequencing using the Oxford Nanopore Technologies sequencing platform (BioProject ID-PRJNA118816).

Following the generation and sequencing of the recombinant viruses, their characterization was performed in Vero E6 (ATCC® CRL-1586™) and Vero E6 TMPRS2 (JCRB Cell Bank, JCRB1819) cells. While their growth kinetics were performed in Vero E6, Vero E6 TMPRSS2, and Calu-3 (ATCC® HTB-55™) cell lines using the same protocol described by Nooruzzaman et al. (Nooruzzaman et al., 2024).

The growth kinetics of the recombinant viruses were studied over 48 hours in Vero E6, Vero E6 TMPRSS2, and Calu-3 cells. Cells were seeded in 12-well plates at a density of 1.2 × 10□ cells/mL and incubated for 24 hours until they reached 80-90% confluence. The cells were then infected (MOI of 0.1) with the recombinant wild-type (rWA1 and rWA1-nsp1-BA.5.2.1) and recombinant nsp1 variant viruses (rWA1-nsp1-BA.5.2.1Δ82-85 and rWA1-nsp1-BA.5.2.1Δ83-86) and incubated at 4°C for 1 hour to allow virus adsorption. Following adsorption, the inoculum was replaced with 1 mL of complete growth medium, and the plates were incubated at 37°C. Cells and supernatants were collected every 4 hours up to 48 hours post-inoculation and stored at -80°C. The time point “0” represented an aliquot of the virus inoculum stored at -80°C immediately after inoculation was completed. Virus titers at each time point were determined in Vero E6 TMPRSS2 cells using end-point dilutions in a 96-well plate and were expressed as TCID□ □/mL, calculated using the Spearman-Karber method as described previously (Martins et al., 2024).

For plaque assay, 3 × 10□ cells/well of Vero E6 and Vero E6 TMPRSS2 were seeded in 6-well plates and cultured for 24 hours until reaching 90% confluency. Cells were then inoculated with SARS-CoV-2 recombinant viruses (10 plaque-forming units/well for Vero E6 TMPRSS2 cells and 30 plaque-forming units/well for Vero E6 cells) and incubated at 37°C for 1 hour. After incubation, the inoculum was removed, and 2 mL of media containing 2X complete growth medium and 0.5% agarose (final concentration: 1X medium with 0.25% agarose) was added to each well. Once the agarose solidified, the plates were transferred to the incubator and incubated at 37°C for 72 hours. Following incubation, the agarose overlay was removed, and cells were fixed with 3.7% formaldehyde for 30 minutes, then stained with 0.5% crystal violet solution for 15 minutes at room temperature. Lastly, the plaque sizes were quantified using Image J Software.

### Luciferase reporter assays using recombinant SARS-CoV-2

We investigated the effect of nsp1 mutations on modulation of the interferon responses using a luciferase assay in the context of SARS-CoV-2 infection in HEK293T-hACE2 cell lines. A luciferase assay was performed using recombinant WT (rWA1 and rWA1-nsp1-BA.5.2.1) and variant viruses (rWA1-nsp1-BA.5.2.1Δ82-85 and rWA1-nsp1-BA.5.2.1Δ83-86) in HEK293T-hACE2 cells. Cells were seeded at a density of 2 × 10□ cells/mL in complete DMEM supplemented with 10% FBS in a 24-well plate, with 0.5 mL per well, and incubated at 37°C in 5% CO□. After 24 hours, the medium was replaced with complete MEM containing 10% FBS, and the plate was returned to the incubator. A transfection mixture containing the plasmids pIFNβ/pNF-κB (200 ng/well) and pRL-TK (50 ng/well) was prepared and incubated with the cells for 6 hours. Following transfection, the cells were infected with the recombinant viruses at an MOI of 3 and incubated for 12 hours. After this incubation period, the cells were stimulated with Sendai virus (SV) (100 HAU, Cantell strain) and TNF-α (25 ng/ml) (Cell Signaling Technology) for 8 hours. The cells were then harvested and resuspended in 1X passive lysis buffer (Promega, USA), and the luciferase assay was performed as described above (Nooruzzaman et al., 2024).

### Pathogenesis studies in hamsters

A total of thirty 60-day-old LVG golden Syrian hamsters (strain 049; 15 female and 15 male) were purchased from Charles River Laboratories, United States. The average body weight of the hamsters was 100.89 g, ranging from 82 to 115 g. All animals were housed in an Animal Biosafety Level 3 (ABSL-3) facility at the East Campus Research Facility (ECRF) at Cornell University. On day 0, the hamsters were inoculated intranasally with 100 µL of recombinant wild-type (rWA1 and rWA1-nsp1-BA.5.2.1) or variant virus (rWA1-nsp1-BA.5.2.1Δ82-85 and rWA1-nsp1-BA.5.2.1Δ83-86) suspension containing 5 × 10□ PFU (n = 6/virus group, 3 males and 3 females) and housed individually in cages. Animals were monitored daily for clinical signs and body weight changes over a 5-day experimental period. Oropharyngeal swabs were collected daily from days 0 to 5, placed in sterile microcentrifuge tubes containing 1 mL of viral transport medium (VTM) (Corning®, Glendale, AZ, USA), and stored at -80°C until further analysis. Half of the hamsters from each group were euthanized on day 3 post-inoculation, and the remaining hamsters were euthanized on day 5 post-inoculation. All animals were humanely euthanized, and nasal turbinate’s, trachea, and lungs were harvested for virological studies. The study procedures were reviewed and approved by the Institutional Animal Care and Use Committee at Cornell University (IACUC approval number 2020-0064).

### Nucleic acid isolation and real-time reverse transcriptase PCR (rRT-PCR)

Nucleic acid was extracted from oropharyngeal swabs (OPS) and tissues collected from the hamsters. A 10% (w/v) homogenate was prepared from nasal turbinate, trachea, and lung tissues in plain DMEM using a stomacher (60-second speed cycle, Stomacher® 80 Biomaster). The tissue homogenate was clarified by centrifugation at 2000 × g for 10 minutes. For RNA extraction, 200 µL of oropharyngeal swab samples and clarified tissue homogenate were processed using the MagMax Core extraction kit (ThermoFisher Scientific, USA) with the automated KingFisher Flex nucleic acid extractor (ThermoFisher Scientific, USA). Subsequently, Real-time RT-PCR (rRT-PCR) was performed using the same protocol described by Nooruzzaman et al. (Nooruzzaman et al., 2024). Relative viral genome copy numbers were calculated using a standard curve and analysed with GraphPad Prism version 9.0.1 (GraphPad, La Jolla, CA, USA).

### Virus isolation and titration

All OPS and tissue homogenates were subjected to virus isolation in Vero E6 TMPRSS2 cells. For virus titration, serial 10-fold dilutions of the samples were prepared in plain DMEM and inoculated into Vero E6 TMPRSS2 cells in 96-well plates. Two days later, the culture supernatant was removed, and the cells were fixed with 3.7% formaldehyde solution before undergoing immunofluorescence analysis (IFA). A mouse monoclonal antibody targeting the SARS-CoV-2 N protein (SARS-CoV-2 anti-N mAb clone B61G11) was used as a primary antibody in the IFA. Virus titers at each time point were determined using end-point dilutions along with the Spearman and Karber methods, and results were expressed as TCID_50_ per mL.

### Statistical analysis

Statistical analysis was performed by 2-way analysis of variance (ANOVA) followed by multiple comparisons. The Mann-Whitney U test was used to compare plaque sizes between the recombinant wild type and variant viruses. Statistical analysis and data plotting were performed using the GraphPad Prism software version 9.0.1 (GraphPad, La Jolla, CA, USA).

## Results

### Computational-assisted structural analysis

To evaluate the sequence of SARS-CoV-2 wild-type and variant nsp1 we performed multiple sequence alignment. The analysis revealed nucleotide deletions between nucleotide position G245 to A253 and C247 to T258 at the N-terminal of the nsp1 protein of SARS-CoV-2 BA.5.2.1 isolate NYI20/22. These deletions led to loss of four-amino-acid in both nsp1 variants, which were defined as nsp1-BA.5.2.1Δ82-85 and nsp1-BA.5.2.1Δ83-86 variants. Analysis of the Global Initiative on Sharing All Influenza Data (GISAID) database revealed that the overall frequency of both mutants is <0.5% based on available sequences as of January, 24 2024.

The impact of these mutations on the structure of nsp1 was examined. We modelled both variants (nsp1Δ82-85 and nsp1Δ83-86) using the Swiss Modeller tool, with the nsp1 crystal structure 8A55 serving as a template. Due to high sequence homology, the variants aligned precisely with the template. Importantly, the Δ82-85 and Δ83-86 deletions in the nsp1 sequences led to the complete loss of a helix and a shortening of the loop (Fig. 1). These models were validated using a Ramachandran plot via the PROCHECK server which showed that 95.8% and 92.6% of the residues fell within the most favoured regions for the nsp1-BA.5.2.1Δ82-85 and nsp1-BA.5.2.1Δ83-86 variants, respectively. The models were further assessed by calculating their RMSD value using PyMOL. The variant showed RMSD values of 0.059 and 0.052 for nsp1-BA.5.2.1Δ82-85 and nsp1-BA.5.2.1Δ83-86 variants, respectively (Fig. 1). Interestingly, the electrostatic surface potential in the deleted region was remarkably altered (Fig. 1). As shown in Fig. 1, both mutants show a noticeable shift in surface potential, with an increase in negative potential in residues around the deleted regions. This shift highlights the significant impact of the deleted amino acids on the nsp1 protein electrostatic environment, which could impact its functional properties and interactions.

**Fig. 1:**
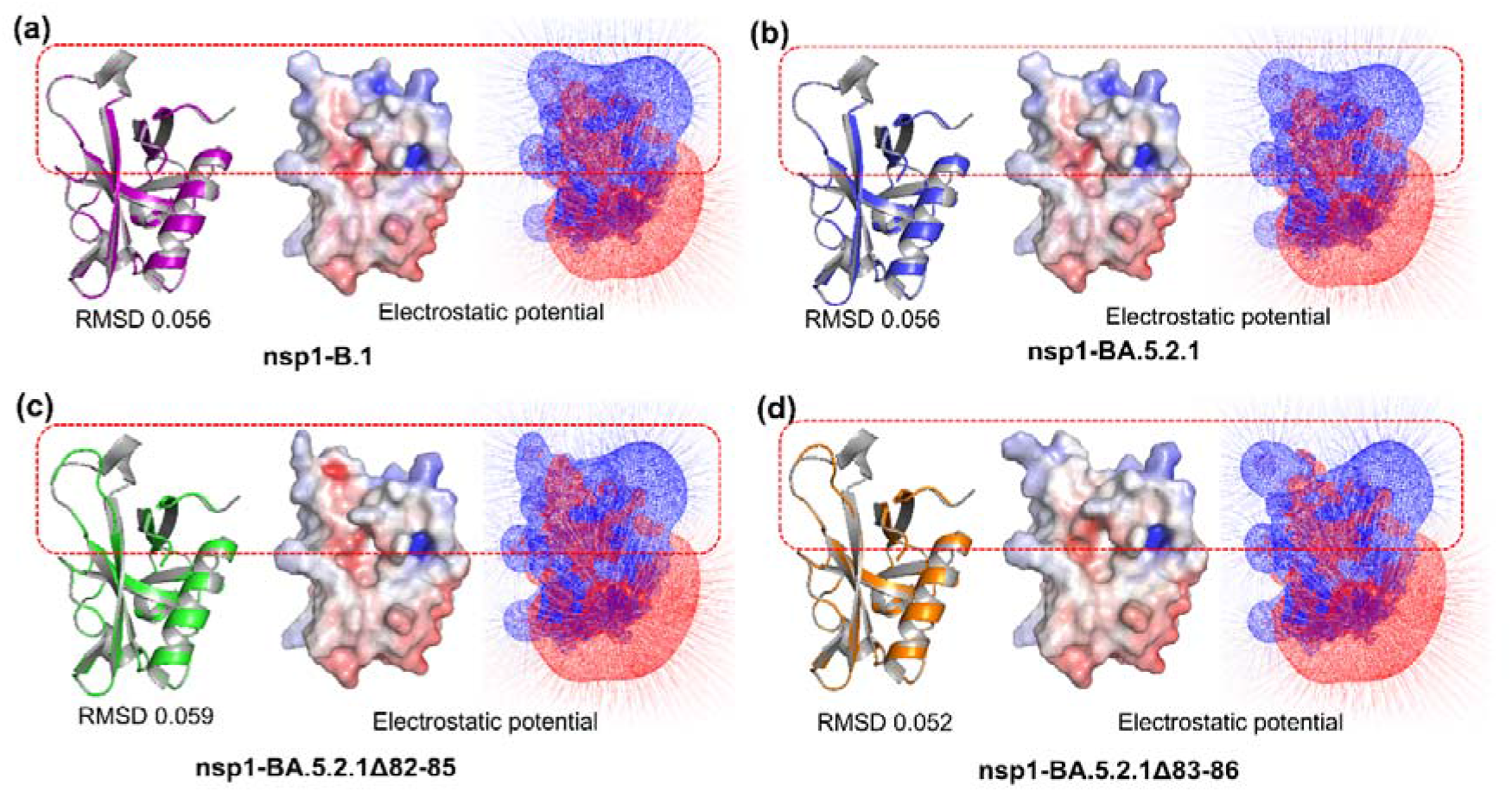
Comparison of wild-type and deleted variants nsp1 proteins. (**a**) nsp1-B.1, (**b**) nsp1-BA.5.2.1, (**c**) nsp1-BA.5.2.1Δ82-85 and (**d**) nsp1-BA.5.2.1Δ82-85. (a) Cartoon representation of the superimposed structures of the template protein (grey) and the nsp1-B.1 variant (purple), highlighting the specific deletion with a red dotted rectangle. Next, surface representation of the electrostatic potential of nsp1-B.1, with negative potential regions displayed in red and positive potential regions in blue. Electrostatic potential map, showing the mesh and gradient distribution of nsp1-B.1, illustrating the variation in electrostatic potential across the protein surface. (b) Cartoon representation of the superimposed structures of the template protein and the nsp1-BA.5.2.1 variant (blue), highlighting the specific deletion with a red dotted rectangle. Surface representation of the electrostatic potential of nsp1-BA.5.2.1, with negative potential regions displayed in red and positive potential regions in blue. Electrostatic potential map, showing the mesh and gradient distribution of nsp1-BA.5.2.1, illustrating the variation in electrostatic potential across the protein surface. (c) Cartoon representation of the superimposed structures of the template protein (grey) and the nsp1-BA.5.2.1Δ82-85 variant (green), highlighting the specific deletion with a red dotted rectangle. Surface representation of the electrostatic potential of nsp1-BA.5.2.1Δ82-85, with negative potential regions displayed in red and positive potential regions in blue. Electrostatic potential map, showing the mesh and gradient distribution of nsp1-BA.5.2.1Δ82-85, illustrating the variation in electrostatic potential across the protein surface. (d) Cartoon representation of the superimposed structures of the template protein (grey) and the nsp1-BA.5.2.1Δ83-86 variant (orange), highlighting the specific deletion with a red dotted rectangle. Surface representation of the electrostatic potential of nsp1-BA.5.2.1Δ83-86, with negative potential regions displayed in red and positive potential regions in blue. Electrostatic potential map, showing the mesh and gradient distribution of nsp1-BA.5.2.1Δ3-86, illustrating the variation in electrostatic potential across the protein surface. The proteins models were generated using Swiss MODELLER, resulting in RMSD values of 0.059 Å for nsp1-BA.5.2.1Δ82-85 and 0.052 Å for nsp1-BA.5.2.1Δ83-86, indicating high structural similarity.

### Luciferase reporter activity assay for evaluating nsp1 function

The nsp1 protein of SARS-CoV-2 plays a crucial role in modulating the host immune response by downregulating the expression of ISGs, which are vital for the host’s antiviral defense. In this study, we investigated whether the nsp1 variants (nsp1-BA.5.2.1Δ82-85 and nsp1-BA.5.2.1Δ3-86) of SARS-CoV-2 affect downstream signaling in the interferon cascade (Fig. 2). We assessed the effect of the wild-type (nsp1-B.1 and nsp1-BA.5.2.1) and variants nsp1 (nsp1-BA.5.2.1Δ82-85 and nsp1-BA.5.2.1Δ3-86) proteins on IFN inhibition. Our results showed that the expression of wild-type nsp1 (nsp1-B.1 and nsp1-BA.5.2.1) inhibited NF-κB, IFN-β and IRF3 promoter activation. In contrast, the nsp1 variants (nsp1-BA.5.2.1Δ82-85 and nsp1-BA.5.2.1Δ3-86) exhibited significantly reduced inhibitory activity (Fig. 2). Moreover, WT nsp1 (nsp1-B.1 and nsp1-BA.5.2.1) effectively reduced SeV-induced activation of IFN-β, NF-κB and IRF3, while the inhibition was not observed in cells transfected with the variant (nsp1-BA.5.2.1Δ82-85 and nsp1-BA.5.2.1Δ3-86) nsp1 proteins (Fig. 2). Therefore, our findings indicate that the amino acid residues from 82–85 and 83–86 in the N-terminus of nsp1 are critical for its inhibitory function on IFN signalling.

**Fig. 2:**
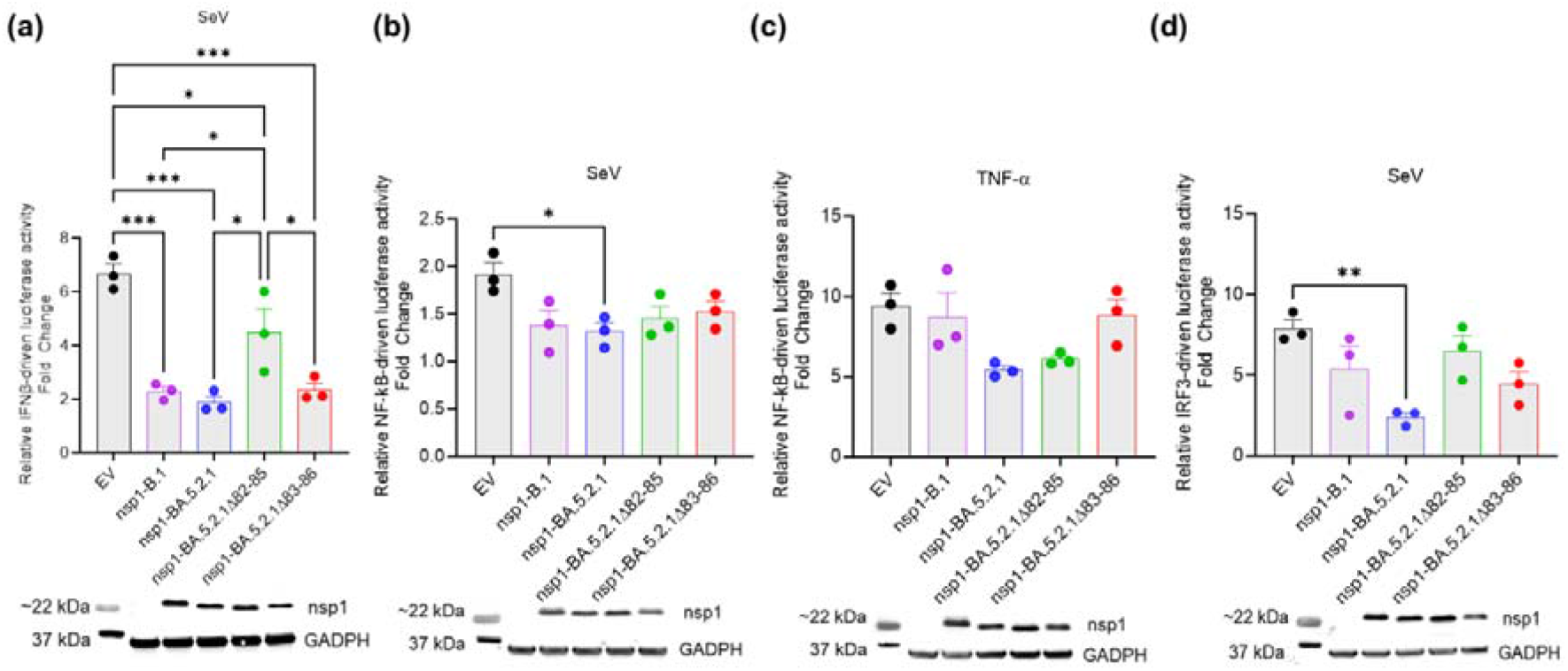
Quantification of luciferase activity in HEK293T cells transfected with nsp1 WT and variant with/without SEV/TNF-α stimulation. Data indicates mean ± SEM of three independent experiments. One-way analysis of variance with multiple comparison test, *P□< □0.05, **P□< □0.01, and ***P□< □0.001. Protein expressions were detected by Western blot and represented as similar manner as shown in luciferase activity assay. GADPH was used as a loading control. All experiments were done repeated three times as independent experiment, and one representative of experimental blot is shown. EV: Empty Vector.

### Replication properties of rSARS-CoV-2 virus

The replication kinetics, plaque size, and morphology of the rSARS-CoV-2 variants were investigated and compared to rWA1, rWA1-nsp1-BA.5.2.1 and nsp1 variants (rWA1-nsp1-BA.5.2.1Δ82-85 and nsp1-BA.5.2.1Δ3-86) *in vitro* (Fig. 3a). Both recombinant nsp1 variants (rWA1-nsp1-BA.5.2.1Δ82-85 and nsp1-BA.5.2.1Δ3-86) exhibited similar replication kinetics in Vero E6, Vero E6 TMPRSS2, and Calu-3 cells, demonstrating that the recombinant WTs (rWA1, rWA1-nsp1-BA.5.2.1) and variants viruses (rWA1-nsp1-BA.5.2.1Δ82-85 and rWA1-nsp1-BA.5.2.1Δ83-86) present similar replication properties in these cells (Fig. 3b). Additionally, while the rWA1-nsp1-BA.5.2.1 and nsp1 variants (rWA1-nsp1-BA.5.2.1Δ82-85 and rWA1-nsp1-BA.5.2.1Δ83-86) produced comparable plaques (sizes) in Vero E6 TMPRSS2 cells, their plaques were smaller than those produced by rWA1 (****P□≤ □0.0001) (Fig. 3c).

**Fig. 3:**
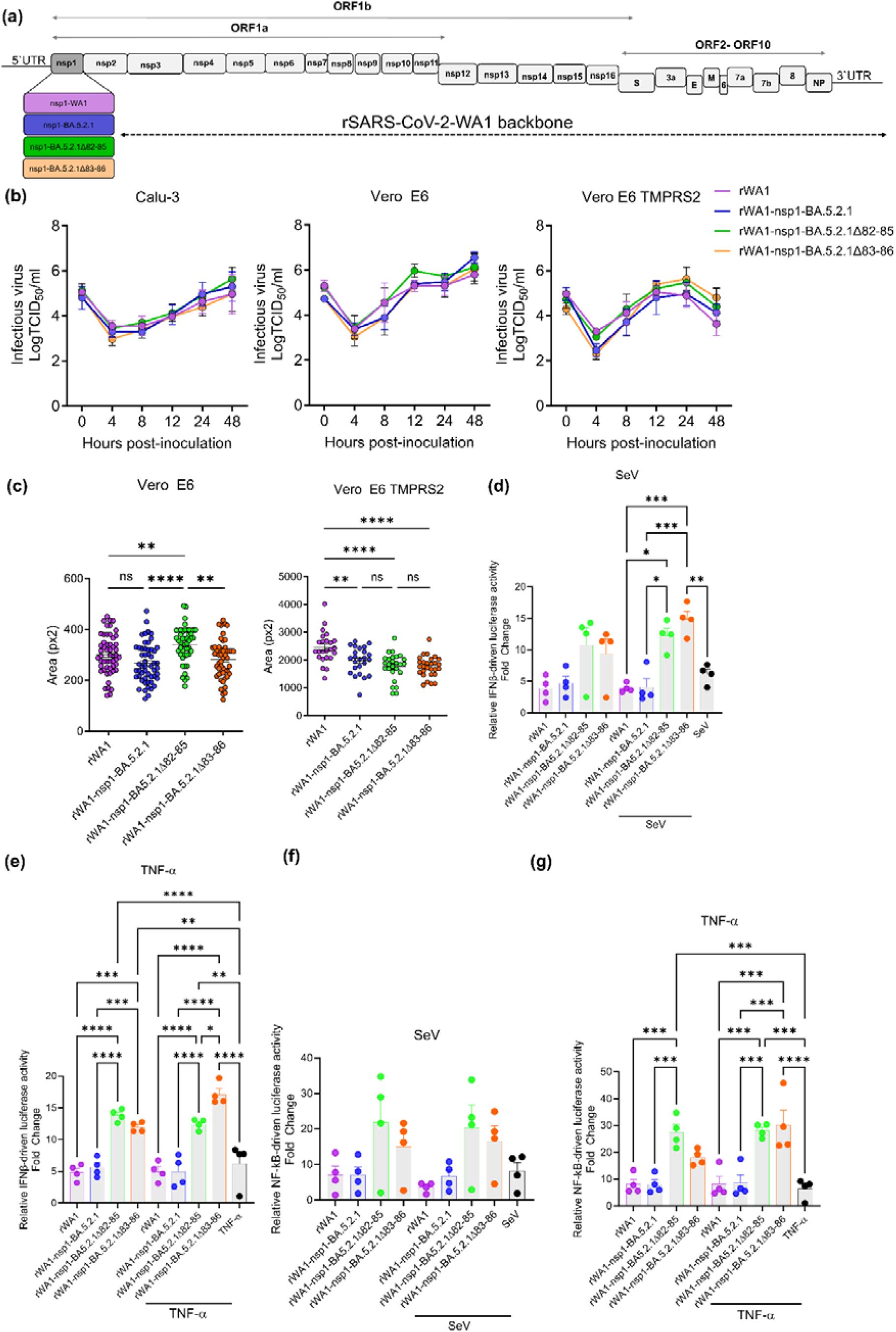
Characterization of recombinant WTs (rWA1 and rWA1-nsp1-BA.5.2.1) and variant viruses (rWA1-nsp1-BA.5.2.1Δ82-85 and nsp1-BA.5.2.1Δ3-86). (a) A schematic representation of the rSARS-CoV-2 genome where the nsp1 of rWA1 is replaced by nsp1-BA.5.2.1 and its variants in the recombinant virus. The locations of other viral proteins and untranslated regions (5□ UTR and 3□ UTR) are shown for WA1 only. (b) The growth kinetics of recombinant WTs (rWA1 and rWA1-nsp1-BA.5.2.1) and variant viruses (rWA1-nsp1-BA.5.2.1Δ82-85 and nsp1-BA.5.2.1Δ3-86) were assessed in Vero E6, Vero E6 TMPRSS2, and Calu-3 cells. Cells were infected at a multiplicity of infection (MOI) of 0.1. At 4-, 8-, 12-, 24-, and 48-hours post-infection, the presence of infectious virus in tissue culture supernatants was determined using TCID□ □/mL, calculated by the Spearman-Karber method. (c) Plaque phenotype of rSARS-CoV-2 was assessed in Vero E6 and Vero E6 TMPRSS2 cells. Cells were infected with recombinant WTs (rWA1 and rWA1-nsp1-BA.5.2.1) and variant viruses (rWA1-nsp1-BA.5.2.1Δ82-85 and nsp1-BA.5.2.1Δ83-86) (∼30 PFU for Vero E6 and ∼10 PFU for Vero E6 TMPRSS2), respectively. At 72 hours post-infection, viral plaques were observed under a microscope and the areas of selected plaques were measured and compared for each rSARS-CoV-2 virus. (d-g) Quantification of luciferase activity in HEK293T-hACE2 cells transfected with rSARS-CoV-2 WT and variant viruses with/without SEV/TNF-α stimulation. n=4, *P□< □0.05, **P□< □0.01, and ***P□< □0.001, one-way analysis of variance with multiple comparison test.

### Mutations in nsp1 lead to decreased inhibitory activity on the IFNβ-signaling pathway during SARS-CoV-2 infection

To determine whether the deletions in nsp1 affected the activation of the IFN and NF-κB signalling pathways, we assessed IFN- and NF-κB-mediated luciferase activity in HEK-293T-hACE2 cells in the context of infection with recombinant WTs (rWA1 and rWA1-nsp1-BA.5.2.1) and variant (rWA1-nsp1-BA.5.2.1Δ82-85 and rWA1-nsp1-BA.5.2.1Δ83-86) viruses. While infection with WT viruses (rWA1 and rWA1-nsp1-BA.5.2.1) efficiently blocked SEV and TNFα induced activation of IFNβ and NF-κB-driven luciferase activity, infection with variant (rWA1-nsp1-BA.5.2.1Δ82-85 and rWA1-nsp1-BA.5.2.1Δ83-86) did not efficiently prevent activation of these signalling pathways as evidenced by increased luciferase activity in cells infected with the variant viruses when compared to WT virus infected cells (Fig. 3d-g). These results suggest that the deletions in the nsp1 significantly impair its ability to modulate these critical antivirals and pro-inflammatory signaling pathways (Fig. 3d-g).

### Deletions in nsp1 do not affect pathogenesis in a hamster model

To determine the role of nsp1 deletions to SARS-CoV-2 virulence and disease, we conducted a pathogenesis study in hamsters (Fig. 4a). While control animals gain weight throughout the 5-day experimental period, all inoculated animals lost weight until day 3 pi (Fig. 4b). After day 3 pi most inoculated animals except for those infected with rWA1-nsp1-BA.5.2.1Δ82-85, gain weight until the end of the experiment (Fig. 4b). Viral RNA load, as determined by rRT-PCR in OPS samples collected from days 0 to 5 pi, revealed similar levels of viral RNA for both rWTs (rWA1 and rWA1-nsp1-BA.5.2.1) and the variant (rWA1-nsp1-BA.5.2.1Δ82-85 and rSARS-CoV-2-nsp1-BA.5.2.1Δ83-86) viruses (Fig. 4c).

**Fig. 4.**
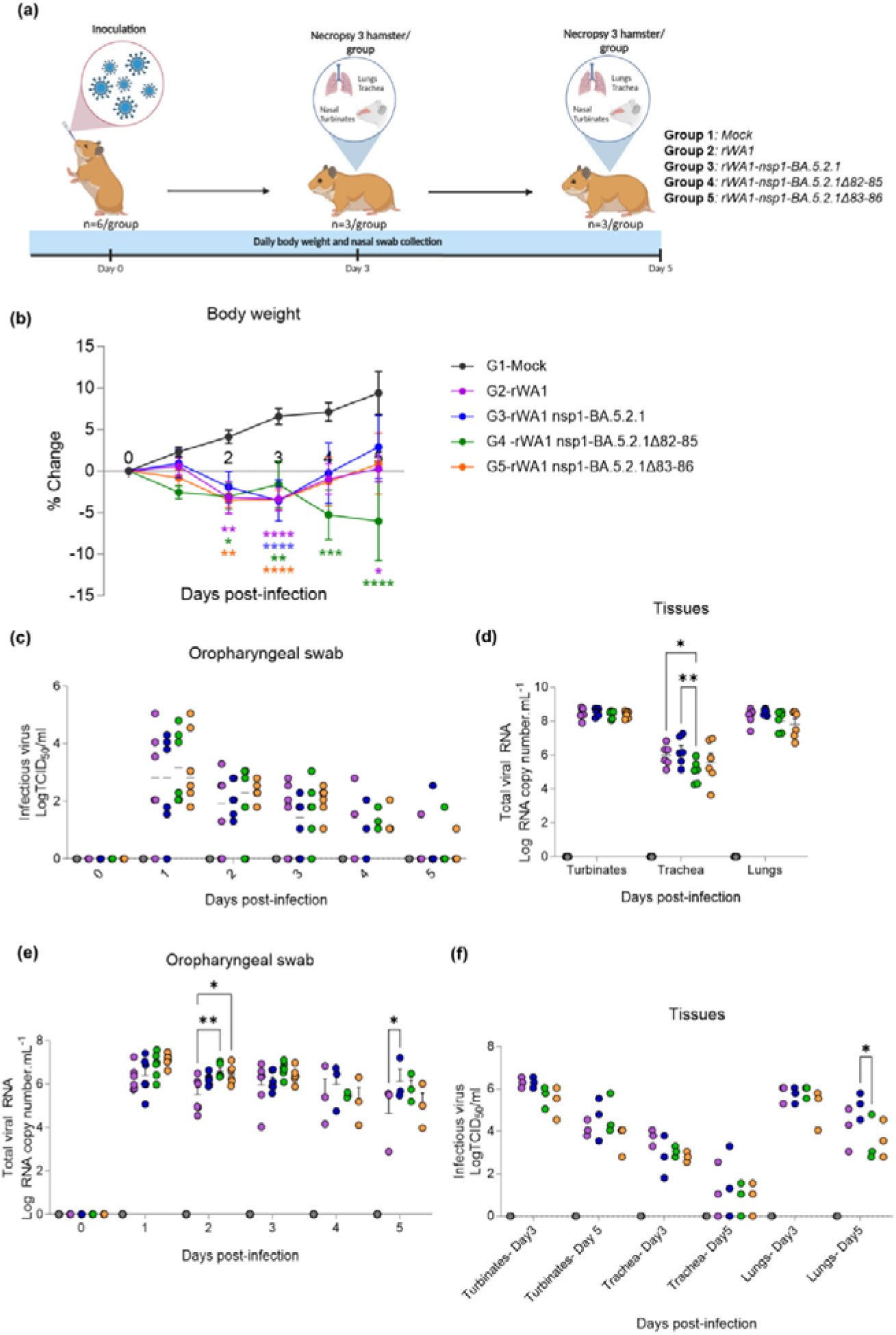
Pathogenesis study in hamsters. (a) Experimental design animal pathogenesis study in a hamster model of SARS-CoV-2 infection. Animals were allocated into five groups and monitored over a 5-day experimental period. (b) Changes in body weight of hamsters following intranasal inoculation with recombinant WTs (rWA1 and rWA1-nsp1-B5.2.1) and variant (rWA1-nsp1-BA.5.2.1Δ82-85 and rWA1-nsp1-BA.5.2.1Δ83-86) viruses over the 5-day period. Viral RNA load in oropharyngeal swabs (c) and tissues (d) quantified by rRT-PCR. Data represent mean□± □SEM of 3-6 animals per hamster/group/day. (e) Infectious viral loads in oropharyngeal swabs determined using endpoint titrations and expressed as TCID50.mL^−1^. (f) Viral RNA load in the nasal turbinate, trachea, and lungs, quantified by rRT-PCR.

Viral load in tissues were assessed on days 3 and 5 pi, in nasal turbinates, trachea, and lungs samples. No significant differences in viral RNA load in nasal turbinate and lungs were observed between rWTs (rWA1 and rWA1-nsp1-BA.5.2.1) and variant (rWA1-nsp1-BA.5.2.1Δ82-85 and rWA1-nsp1-BA.5.2.1Δ83-86) viruses inoculated hamsters. A significant difference in viral RNA load was observed in the trachea, where rWA1-nsp1-BA.5.2.1Δ82-85 exhibited a lower viral level when compared to both WTs (Fig. 4d). Additionally, the infectious viral load detected in both OPS and tissue samples (turbinates, trachea, and lungs) displayed no significant variation between WTs (rWA1 and rWA1-nsp1-BA.5.2.1) and variant (rWA1-nsp1-BA.5.2.1Δ82-85 and rWA1-nsp1-BA.5.2.1Δ83-86) viruses (Fig. 4e-f). Viral titers in OPS, decreased progressively in all infected groups, consistent with their recovery (Fig. 4e). While viral loads in nasal turbinates and lungs were comparable between both recombinant wild types (rWA1 and rWA1-nsp1-BA.5.2.1) and variants, the trachea showed a lower viral burden in all infected groups (Fig. 4f). Importantly, no major changes in viral load between the groups were observed. These results demonstrate that nsp1 mutations in the N-terminus (Δ82-85 and Δ83-86) do not markedly affect SARS-CoV-2 virulence and pathogenesis in the hamster model.

## Discussion

The nsp1 of SARS-CoV-2 has evolved diverse mechanisms and functions to manipulate the host translation machinery while allowing efficient translation of viral mRNAs (Kamitani et al., 2006; Thoms et al., 2020). In this study, we identified two distinct deletion mutations in the N-terminal region of nsp1 (nsp1Δ82-85 and nsp1Δ83-86) and examined their structural and functional impacts in comparison with WT-nsp1.

Structural analysis of WT and variant nsp1 proteins revealed that the deletion mutations in the nsp1 significantly altered the protein’s structural pocket. This disruption also caused changes in its electrostatic potential, with the nsp1 variants showing a more negative electrostatic potential compared to the region where the amino acids were deleted. These findings suggest that the deleted amino acids could have significant implications for its functional properties.

The SARS-CoV-2 nsp1 protein is known to interfere with interferon signalling, a critical component of the host’s antiviral response (Fisher et al., 2022; Thoms et al., 2020). However, using a luciferase assay, we found that these nsp1 variants did not inhibit interferon signalling, indicating a functional impairment caused by the deletions. We also assessed the effect of these mutations on viral replication. Interestingly, despite their altered impact on interferon signalling, the nsp1 mutations did not significantly affect viral replication properties in three different cell lines or in the hamster model during the experimental infection. This suggests that the variants retain similar viral replication efficiency to the WT virus across different cell types. Our results also align with previous studies by Lin et al., 2021, who also reported nsp1 mutations that affect interferon signalling. Their findings support our conclusion that certain mutations in N-terminal region of nsp1 can disrupt its role in immune evasion without significantly affecting viral replication and pathogenesis. These findings lead us to conclude that the four amino acid deletions in the nsp1 variants specifically impair their ability to disrupt interferon signalling. One limitation of the approach we used to generate the recombinant viruses containing the nsp1 Δ82-85 and Δ83-86 mutations is the use of the WA1 virus backbone. Since these natural mutations were identified in the Omicron BA5.2.1 variant, it would be important to evaluate their effect in the context of an Omicron BA5.2.1 background.

In conclusion, our study demonstrates that the deleted four amino acids in the N-terminal region of SARS-CoV-2 nsp1 play a crucial role in the inhibition of interferon signalling as these deleted amino acids revealed the intricate interplay between viral proteins and the host innate immune response. These findings enhance our understanding of SARS-CoV-2 pathogenesis and provide insights that aimed to completely understand the mechanism of action of the deleted amino acid of nsp1 on interferon signalling.

## Acknowledgements

We thank the Center for Animal Resources and Education (CARE) staff and Cornell Biosafety team for the support. This work was funded in part by the National Institutes of Health (NIH) and National Institute of Allergy and Infectious Diseases (NIAID) (grant no. R01AI166791-01).

